# APOE*4 genotype, blood-brain barrier leakage, ischaemic stroke subtype and location*

**DOI:** 10.64898/2026.06.05.730537

**Authors:** Krystal K. Laing, Maria del C. Valdés Hernández, J Michael, Thrippleton, Stephen Makin, Francesca M. Chappell, Owen Dando, Deepali Vasoya, Paul Armitage, Joanna M. Wardlaw

## Abstract

**Background:** Apolipoprotein E (APOE) has been implicated in blood–brain barrier (BBB) dysfunction and may influence ischaemic cerebrovascular disease and cerebral small-vessel disease (cSVD). This study examined associations between APOE genotype, BBB permeability, and infarct distribution in patients with mild ischemic stroke.

**Methods:** We recruited patients with mild ischemic stroke who underwent structural and dynamic contrast-enhanced MRI (DCE-MRI) and APOE genotyping. Infarct type and location, white matter hyperintensities (WMH), and perivascular spaces (PVS) were assessed. BBB-related metrics were quantified using fractional plasma volume (vP) and permeability–surface area product (PS) across five brain regions of interest: deep grey matter (DGM), hippocampus, thalamus, normal-appearing white matter (NAWM), and WMH. Associations between genotype, BBB metrics, vascular risk factors, and age were evaluated using linear mixed-effects models. Binary logistic regression was also applied to assess the association between APOE status and infarct location by vascular territory (anterior vs posterior circulation).

**Results:** Among 147 patients with APOE genotype and BBB measures, APOE4 carriers (n=44) demonstrated a greater proportion of posterior circulation infarcts than E3/E3 individuals (n=80; 56.4% versus 32.9%), including higher frequencies of posterior cerebral artery cortical, posterior borderzone, and thalamic infarcts. Mean PS and vP did not differ significantly by genotype. Increasing age was associated with lower PS across multiple regions and lower vP in WMH, while higher vascular risk burden was associated with lower vP in NAWM and WMH. Inclusion of regional BBB metrics did not substantially alter APOE4 effect estimates in infarct-location models.

**Conclusions:** APOE4 carriers showed a posterior-predominant infarct distribution despite similar BBB permeability and vascularity measures. Age and vascular risk burden were more strongly associated with BBB-related imaging metrics than APOE genotype. These findings add to evidence suggesting that APOE genotype may influence regional cerebrovascular vulnerability and that this effect is unlikely to be fully explained by DCE-MRI–derived measures of BBB permeability and vascularity alone.

## 1. Introduction

Apolipoprotein E (APOE) is a multifunctional lipid transporter involved in lipid metabolism. Its three major isoforms – E2, E3, E4 – differ by amino acid substitutions at residues 112 and 158, resulting in isoform-dependent variability in circulating levels of cholesterol^1^. The APOE*4* variant is associated with elevated plasma total and low-density lipoprotein (LDL) cholesterol, contributing to atherosclerosis^2,3^ and representing a well-established, although incompletely understood, genetic risk factor for Alzheimer’s Disease ^4,5^.

Prior meta-analyses^2,6,7^ have identified the E4 allele as a potential genetic marker for ischaemic stroke, though subsequent studies reported inconsistent findings^8,9^. A more recent meta-analysis^10^ examining APOE polymorphisms across ischaemic stroke subtypes suggested an association between the E4 and lacunar (intrinsic small vessel) stroke, but not with large artery atheromatous (LAA) or cardioembolic (CE) ischaemic stroke. These findings suggest that APOE-related stroke risk may not be solely mediated through lipid regulation or atheroma consistent with increasing evidence that cerebral small vessel disease (cSVD), is a non-atheromatous intrinsic microvessel abnormality^11,12^.

Recent studies have also reported associations between APOE status and white matter hyperintensities (WMH) location. Cohort studies including Washington Heights–Inwood Columbia Aging Project (WHICAP; New York City, USA) and Étude de Santé Psychologique Prévalence, Risques et Traitement (ESPRIT; Montpellier, France) reported greater WMH volume in parietal and occipital than other brain regions among ε4 carriers^13,14^. Furthermore, analyses comparing APOE*4* carriers and non-carriers have shown that greater WMH volume is associated with worse attention/executive functions, memory, and language performance in E4 carriers, suggesting that APOE-related microvascular changes may contribute to regional brain vulnerability and cognitive decline^15^. Consistent with this, APOE ε4 homozygotes showed reduced white matter integrity at older ages in UK Biobank^14,16^.

cSVD is a disorder of small perforating vessels (arterioles, capillaries, and venules), with effects observable as WMH, lacunes, microbleeds, and perivascular spaces (PVS) on magnetic resonance imaging (MRI) or pathology and impaired integrity of normal appearing tissues. A growing body of literature supports the idea that abnormal microvessel function, including of the blood brain barrier (BBB), may contribute to the cSVD development^17–21^. BBB permeability increases with ageing, and is linked to cognitive impairment, including in AD and vascular dementia (VaD)^21^.

Both clinical and preclinical studies have reported microvascular abnormalities in E4 carriers, including basement membrane thinning, altered cerebral blood flow, and increased BBB permeability, with progression occurring as early as midlife^22–27^. Some studies also suggest that regional alterations in BBB permeability appear to be more pronounced with ageing, particularly in structures such as the hippocampus and cortex^26,28^.

In the present study, we investigated whether APOE genotype is associated with BBB function and ischaemic stroke subtype (lacunar cSVD and LAA/CE), in patients with minor ischaemic stroke. We hypothesised that APOE*4* would be associated with increased BBB leakage and might show associations with infarct type and location, reflecting genotype-related variation in regional vulnerability of the cerebral microvasculature.

## 2. Methods

### 2.1 Patient Recruitment

We prospectively recruited patients with mild (ie expected to be nondisabling in the long term) ischemic stroke (modified Rankin Scale [mRS] ≤2), presenting to the Lothian regional stroke service between May 2010 to May 2012, with 3-year follow-up. Eligible patients were aged ≥18 years, able to provide informed consent, and able to undergo MRI with a gadolinium-containing contrast agent. Patients with haemorrhagic stroke, severe ischaemic stroke, or transient ischemic attack (TIA; symptoms resolving within 24 hours) were excluded.

Recruitment procedures, inclusion criteria, and medical assessment have been described previously^29,30,20,31,21^. Briefly, *‘mild’ ischemic stroke* was defined as a focal onset of neurological symptoms lasting > 24 hours without an alternative explanation for the symptoms, with NIH Stroke Scale (NIHSS) <8 at its worst point, and not expected to result in long-term dependency (mRS ≤2). All patients were assessed by a stroke physician, and an expert panel confirmed stroke diagnosis and subtype based on clinical findings and MRI features. Stroke subtype was classified clinically using the Oxfordshire Community Stroke Project (OCSP) classification^32^. When discrepancies occurred between clinical and imaging classifications, the imaging subtype (lacunar or cortical) was used^33^.

The Lothian Ethics of Medical Research Committee (REC 09/81,101/54) and NHS Lothian R+D Office (2009/W/NEU/14) approved the study. All patients provided written informed consent.

### 2.2 MRI Protocol

MRI was performed on a 1.5T GE Signa Horizon HDxt clinical scanner (General Electric, Milwaukee, WI, USA) using an 8-channel phased array head-coil^21,29^.

Baseline diagnostic sequences included T1-weighted, T2-weighted, fluid-attenuated inversion recovery (FLAIR), gradient-recalled echo (GRE), and diffusion tensor imaging (DTI), obtained at the time of diagnosis.

Most patients were managed through a rapid-access outpatient stroke prevention clinic, as described previously^29,34,35^.

#### 2.2.1 Dynamic contrast-enhanced magnetic resonance imaging

Dynamic contrast-enhanced magnetic resonance imaging (DCE-MRI), has enabled *in vivo* quantification of regional differences in BBB integrity^23,24,36^. DCE-MRI was performed 1-3 months post-stroke to minimize acute effects of infarction on blood-brain barrier (BBB) integrity^20,21^. Two 3D fast-spoiled gradient-echo (FSPGR) acquisitions (flip angles of 2° and 12°) were used to generate pre-contrast T1 (T_10_) maps (TR/TE = 8.24/3.1 ms; field of view 24×24cm, acquisition matrix 256 × 192; 42 slices × 4 mm thickness)^37^.

Gadoterate meglumine (Gd-DOTA, DOTAREM; Guerbet, Paris, France) was administered at 0.2 mL/kg (0.1 mmol/kg body weight) at 2 mL/second via injection pump.

Post-contrast 3D T1w images were acquired 20 times over 24 minutes, achieving a temporal resolution of approximately 73s. The extended acquisition time allowed for detection of subtle BBB leakage^20,21,38^.

### 2.3 Image Processing

MRI data were analysed blind to clinical and APOE data. Structural MRI analyses were conducted independently of DCE data.

Stroke lesions (index [recent acute] and old infarcts), WMH (Fazekas score^39^), lacunes, PVS, microbleeds, and brain atrophy were assessed visually using validated rating scales in accordance with STRIVE recommendations^34^.

The index infarct was classified according to vascular territory (anterior cerebral artery [ACA], middle cerebral artery [MCA], posterior cerebral artery [PCA], and basilar artery [BA]) and anatomical location (cortical, subcortical, or posterior fossa). Lesion size (proportion of vascular territory affected) and presence of swelling were recorded using an extensively validated scale^40^. Patients without a recent infarct visible on MRI were excluded from the infarct localisation analysis.

For computational processing, infarcts were semi-automatically segmented on FLAIR using Analyze 11.0 (AnalyzeDirect, KS, USA)^37^. WMH volumes were quantified from FLAIR images using a previously described segmentation pipeline^29^. Briefly, tissue boundaries and WMH were identified using a multispectral colour fusion technique, which combines pairs of MRI sequences to enhance contrast between normal tissues and lesions.

Deep grey matter (DGM), normal appearing white matter (NAWM), hippocampus, and thalamus masks were generated within this pipeline. Subcortical structures were extracted using FMRIB Software Library (https://fsl.fmrib.ox.ac.uk/fsl/docs/) tools and manually inspected and edited where necessary. To obtain the BBB permeability values in the structures and regions of interest (ROIs), structural MRI was registered to the pre-contrast FSPGR (12°) images using rigid-body registration (FSL-FLIRT)^41^. Transformation matrices were applied to binary masks.

WMH volume was normalised to intracranial volume (ICV) and expressed as percentage ICV (%ICV).

### 2.4 Dynamic contrast-enhanced MRI analysis

Following established procedures, Patlak modelling was applied to five ROIs: WMH, NAWM, DGM, hippocampus, and thalamus^37,42^.

T_1_ maps^43^, and contrast agent concentration, C_i_^44^ were derived.

Vascular input function (VIF) was manually selected in the superior sagittal sinus. Whole-blood contrast concentration *C_b_(t)* was converted to plasma concentration *C_p_(t)*^37^, using individual haematocrit values.

Patlak modelling was used to fit the Gd-DOTA tissue concentration–time curve, *C*_t_*(t),* using validated publicly available in-house software (https://github.com/mjt320/DCE-functions) in MATLAB (MathWorks, Natick, MA, USA). Model outputs included plasma volume fraction (vP; unitless, reported as ×10⁻²) and permeability–surface area product (PS; min⁻¹, reported as ×10⁻⁴).

BBB leakage was reported as PS, instead of Ktrans, as the blood plasma concentrations are considered to meet assumptions of the model and the measurement process is considered to be accurate (as previously shown^21^), as such Ktrans *≈ PS*^45^.

### 2.5 APOE Genotyping

Baseline blood samples were obtained. Genotyping quality control included four duplicate samples. Individuals of non-European ancestry were excluded, as the analytic pipeline was derived from European ancestry-only cohorts.

Genotyping was performed using the Illumina Infinium OmniExpressExome-8 v1.6 SNP array, which assays approximately 960,000 genome-wide and exome-focused single nucleotide polymorphisms (SNPs).

APOE genotype is determined by two SNP variants (rs429358 and rs7412), which define the three major APOE alleles. Depending on the combination of these variants, individuals may carry one of six common genotypes (ε4ε4, ε4ε3, ε4ε2, ε3ε3, ε2ε3, or ε2ε2).

In this dataset, rs7412 (codon 158) was directly measured, whereas rs429358 (codon 112) was imputed from surrounding SNPs within the dataset^46^, using the 1000 Genomes reference panel^47^ (**Supplementary Table 1**). Following a previously established protocol^48^, genotypes were phased using SHAPEIT^49^ and imputed using IMPUTE2^50^.

The resulting APOE genotype frequencies were similar to those reported in other European ancestry datasets, including the UK Biobank^51^, the Lothian Birth Cohort^52^, and CHARGE^53^ (**Table 1**).

**Table 1.** Genotype frequencies compared to other published datasets.

| Genotypes | MSS2 | UK Biobank | CHARGE | Lothian Birth Cohort |
| --- | --- | --- | --- | --- |
|  | N = 147 | N = 488,377 | N = 38,537 | N = 642 |
| $\epsilon 3\epsilon 3$ (%) | 54.4 | 58.2 | 61.9 | 58.6 |
| $\epsilon 2\epsilon 3$ (%) | 15.6 | 12.3 | 12.3 | 12.0 |
| $\epsilon 3\epsilon 4$ (%) | 25.9 | 23.9 | 21.1 | 24.9 |
| $\epsilon 2\epsilon 4$ (%) | 2.7 | 2.6 | 2.3 | 2.2 |
| $\epsilon 4\epsilon 4$ (%) | 1.4 | 2.4 | 1.8 | 2.0 |
| $\epsilon 2\epsilon 2$ (%) | 0 | 0.6 | 0.6 | 0.3 |

APOE4 carrier status was further validated using TaqMan PCR genotyping of rs429358. Concordance between the imputed and PCR-derived E4 carrier status was high (145/147 samples concordant; 98.6% concordance, φ = 0.968; Cohen’s κ = 0.967) (**Supplementary Tables 2 and 3**). One discordant sample was identified (MSSB216), imputed as E3/E4 but classified as E4/E4 by PCR, and was retained as an E4 carrier. One additional sample (MSSB307) yielded an undetermined PCR result and was excluded from genotype-based analyses.

Patients were grouped according to APOE genotype as E4 carriers (ε4/ε4, ε4/ε3, ε4/ε2), E3 homozygotes (ε3/ε3), or E2/E3 heterozygotes.

No E2/E2 individuals were identified in this cohort.

### 2.6 Statistical Analysis

All analyses were performed in R (version 3.6.3; R Foundation for Statistical Computing, Vienna, Austria). Continuous variables were inspected for normality using histograms and Q–Q plots. Normally distributed variables are presented as mean (SD), with 95% confidence intervals (CI). All model assumptions (normality, homoscedasticity, linearity) were verified using diagnostic plots. Regression results are reported as β estimates or odds ratios (OR) with 95% CI.

A vascular risk score (VRS)^54^ was calculated by summing the presence of hypertension, hyperlipidaemia, diabetes mellitus, and current smoking status (coded as 1 if present and 0 if absent), yielding a score ranging from 0 to 4.

Mixed-effects models included participant ID as a random intercept to account for within-subject repeated tissue measurements. APOE genotype was entered as a categorical predictor with E3/E3 as the reference group.

Given the modest sample size and exploratory nature of the analyses, no formal global adjustment for multiple testing was applied.

#### Infarct location and APOE genotype

Associations between APOE genotype and infarct location were examined using binomial logistic regression, adjusting for age and sex. Infarct location was classified as anterior vs posterior circulation. APOE carrier status was also assessed using a count-based model of posterior vs anterior circulation infarcts, adjusted for age and sex.

#### BBB parameters and APOE genotype

Descriptive statistics for PS and vP were first obtained across APOE genotype groups using one-way ANOVA for descriptive purposes.

Associations between APOE genotype and BBB parameters were examined using linear mixed-effects models with PS or vP as the dependent variable. Fixed effects included: APOE genotype, ROIs, age, sex, WMH volume, and VRS, with participant ID included as a random intercept to account for repeated measures across regions within individuals.

An APOE × ROI interaction term was included to test whether genotype effects differed across brain regions. Comparisons between APOE genotype groups and the E3/E3 reference group were performed using estimated marginal means (emmeans), with Dunnett adjustment for multiple testing, accounting for the mixed-effects model and repeated measures.

Binary logistic regression was used to assess the association between APOE4 status and infarct location by vascular territory (posterior vs anterior circulation), adjusting for age and sex. Separate exploratory models examined whether DGM regional vascularity (vP) or BBB permeability (PS) were associated with vascular territory infarct presentation.

## 3. Results

Of the 264 patients enrolled in the full MSS2 cohort^31^, APOE genotype data were available for 152 samples, including four duplicate samples used for quality control. After removal of duplicate samples and exclusion of one individual of non-European ancestry, 147 unique patients remained who completed the 3-year in-person follow-up. Of these, 23 lacked usable DCE-MRI data, leaving 124 patients for the primary BBB analyses. The genotyped sub-cohort (mean age 65.9±11.5 years, 44.9% female) had a similar age distribution to the full MSS2 cohort (66.0±11.6 vs 68.2±12.1 years), with comparable sex distribution (40.9–47.5% vs 37.6%) (**Table 2**). Within this subset, 23 patients were E2/E3 (15.6%), 80 were E3/E3 (54.4%), and 44 were APOE4 carriers (29.9%; E4/E4, E3/E4, or E2/E4).

**Table 2.** Demographics and clinical characteristics of patients by APOE genotype compared to the full MSS2 cohort. VRS: Scale of 0-4. Yes = 1, No = 0, smoker = previous and recent history of smoking. Fazekas scale: 0-6. NIHSS: 0-7.

| Characteristic | E2/E3 | E3/E3 | E4 Carrier | MSS2 Cohort<br>w/o genotyping |
| --- | --- | --- | --- | --- |
|  | *N* = 23 | *N* = 80 | *N* = 44 | *N* = 117 |
| Mean age (SD) | 65.81<br>(11.8) | 66.53<br>(11.9) | 64.69<br>(10.6) | 68.22 (12.10) |
| Sex, female, *n* (%) | 10 (43.5) | 38 (47.5) | 18 (40.9) | 44 (37.6) |
| Stroke subtype, count (%) |  |  |  |  |
| Cortical | 11 (50) | 44 (59.5) | 21 (53.8) | 58 (54.2) |
| Lacunar | 10 (45.5) | 25 (33.8) | 13 (33.3) | 34 (31.8) |
| Cerebellar | 1 (4.55) | 3 (4.05) | 3 (7.69) | 6 (5.61) |
| Brainstem | 0 (0.00) | 2 (2.70) | 2 (5.13) | 9 (8.41) |
| Patients with $\geq 1$ infarct, *n* (%) | 1 (4.35) | 1 (1.25) | 1 (2.27) | 20 (17.1) |
| NIHSS baseline, median | 1 | 1 | 1 | 1 |
| Fazekas Score, median | 4 | 3 | 2 | 3 |
| PVS Score, median |  |  |  |  |
| Basal ganglia | 2 | 1.5 | 1 | 1 |
| Centrum Semiovale | 1 | 2 | 2 | 2 |
| WMH Volume, mean<br>(SD), %ICV | 1.708<br>(1.683) | 1.463<br>(1.545) | 1.151<br>(1.411) | 1.726 (1.750) |
| Mean arterial pressure,<br>median (IQR) | 64.00<br>(57.67 –<br>75.67) | 64.33<br>(52.58 –<br>71.67) | 61.50<br>(56.17 –<br>69.67) | 58.00 (48.33 –<br>68.00) |
| Pulse pressure, median<br>(IQR) | 59.00<br>(50.00 –<br>71.00) | 60.00<br>(49.25 –<br>78.75) | 58.00<br>(40.00 –<br>69.75) | 61.00 (50.00 –<br>76.00) |
| Vascular Risk Score (VRS) | 2.04 | 1.99 | 2.30 | 2.15 |
| Hypertension, yes, n (%) | 17 (73.9) | 58 (72.5) | 33 (84.1) | 79 (67.5) |
| Hyperlipidaemia, yes, n (%) | 10 (43.5) | 49 (61.3) | 32 (72.7) | 70 (59.8) |
| Smoker, yes, n (%) | 16 (69.6) | 45 (56.3) | 27 (61.4) | 89 (76.1) |
| Diabetes, n (%) | 4 (17.4) | 7 (8.75) | 5 (11.4) | 14 (12.0) |

The distribution of infarct type was similar across APOE groups, with cortical infarcts accounting for approximately 50–60% of cases in each subgroup. Lacunar infarcts comprised 45.5% of cases in E2/E3, 33.8% in E3/E3, and 33.3% in E4 carriers (**Table 2**).

WMH volume (%ICV) was slightly lower in the APOE-genotyped sub-cohort compared with the remainder of MSS2 (1.46±1.54 vs 1.73±1.75) but was similar across APOE groups (E2/E3: 1.71±1.68; E3/E3: 1.46±1.55; E4: 1.15±1.41). Fazekas scores and vascular risk factors were also comparable across groups.

### 3.1 Index Infarct Location

E3/E3 individuals showed the broadest distribution of infarct locations, affecting all vascular territories assessed (**Table 3**). The E2/E3 group (n=22 infarcts) demonstrated a more restricted profile, with no infarcts observed in borderzone regions; however, 50% of infarcts (11/22) occurred in PCA, MCA, or ACA cortical territories, and 27.3% (6/22) involved deep structures (18.2% internal/external capsule; 9.1% thalamus).

**Table 3.**
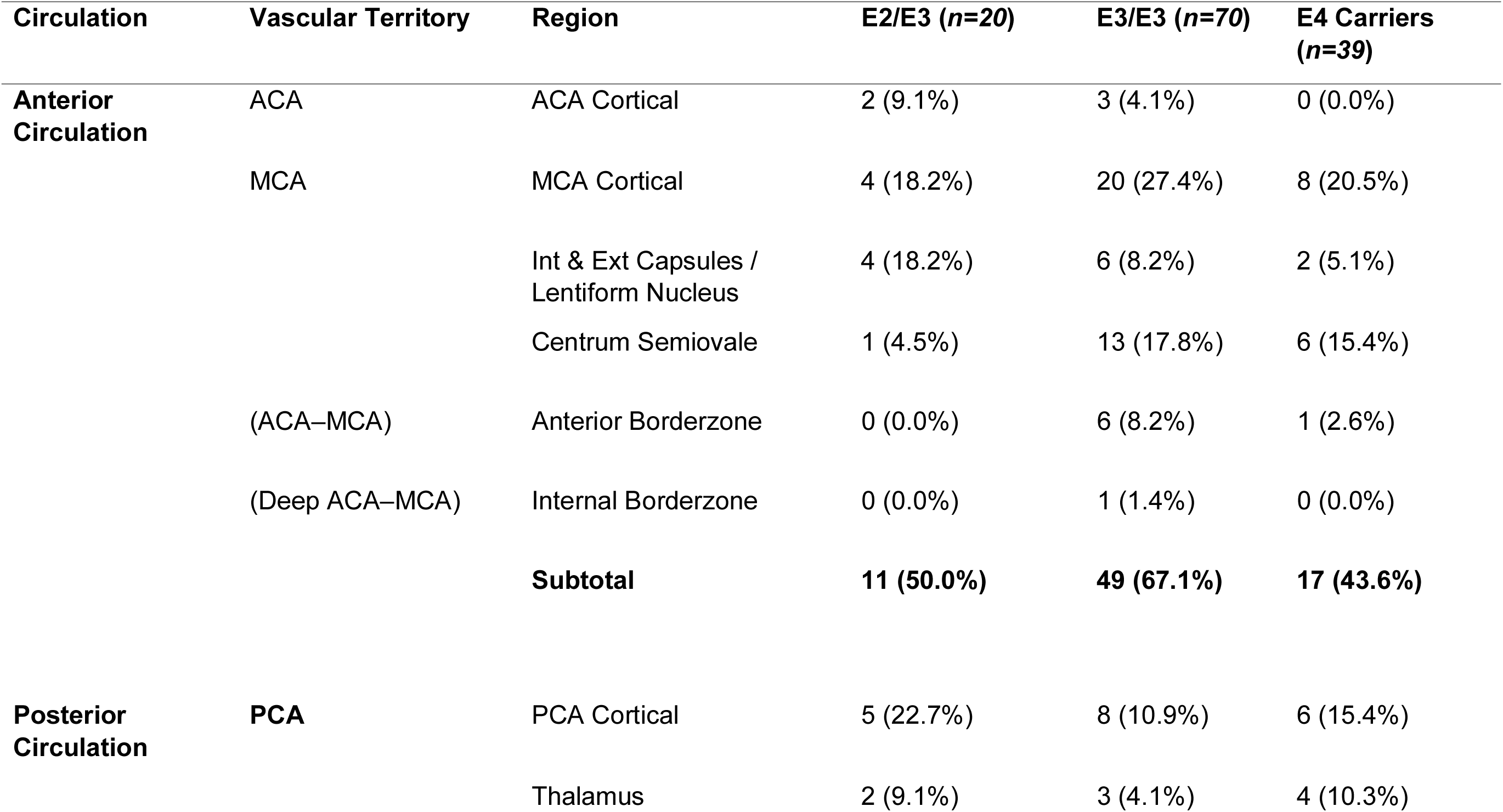
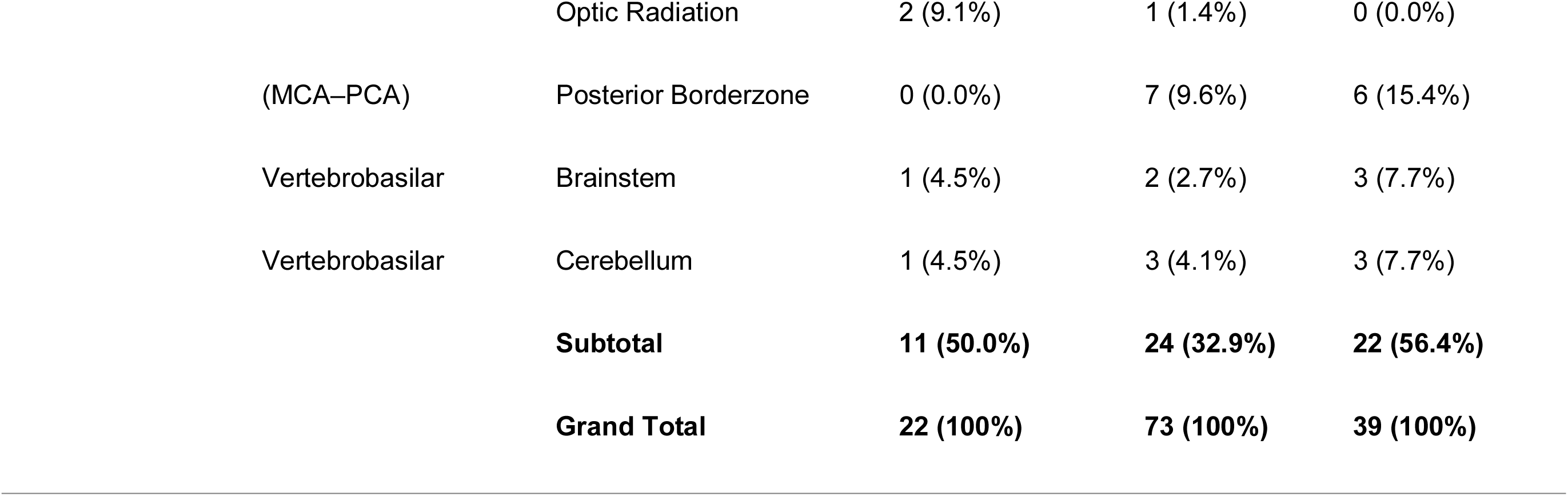
Distribution of index infarcts by vascular territory and APOE genotype. Percentages are based on the number of infarcts within each genotype group. *n indicates the number of patients with visible infarcts*.

E4 carriers showed a higher proportion of infarcts in posterior circulation territories than other APOE groups (**Figure 1**). Compared with E3/E3 individuals, E4 carriers had higher proportions of PCA cortical infarcts (15.4% [6/39] vs 11.0% [8/73]), posterior borderzone infarcts (15.4% [6/39] vs 9.6% [7/73]), and thalamic infarcts (10.3% [4/39] vs 4.1% [3/73]).

**Figure 1.**
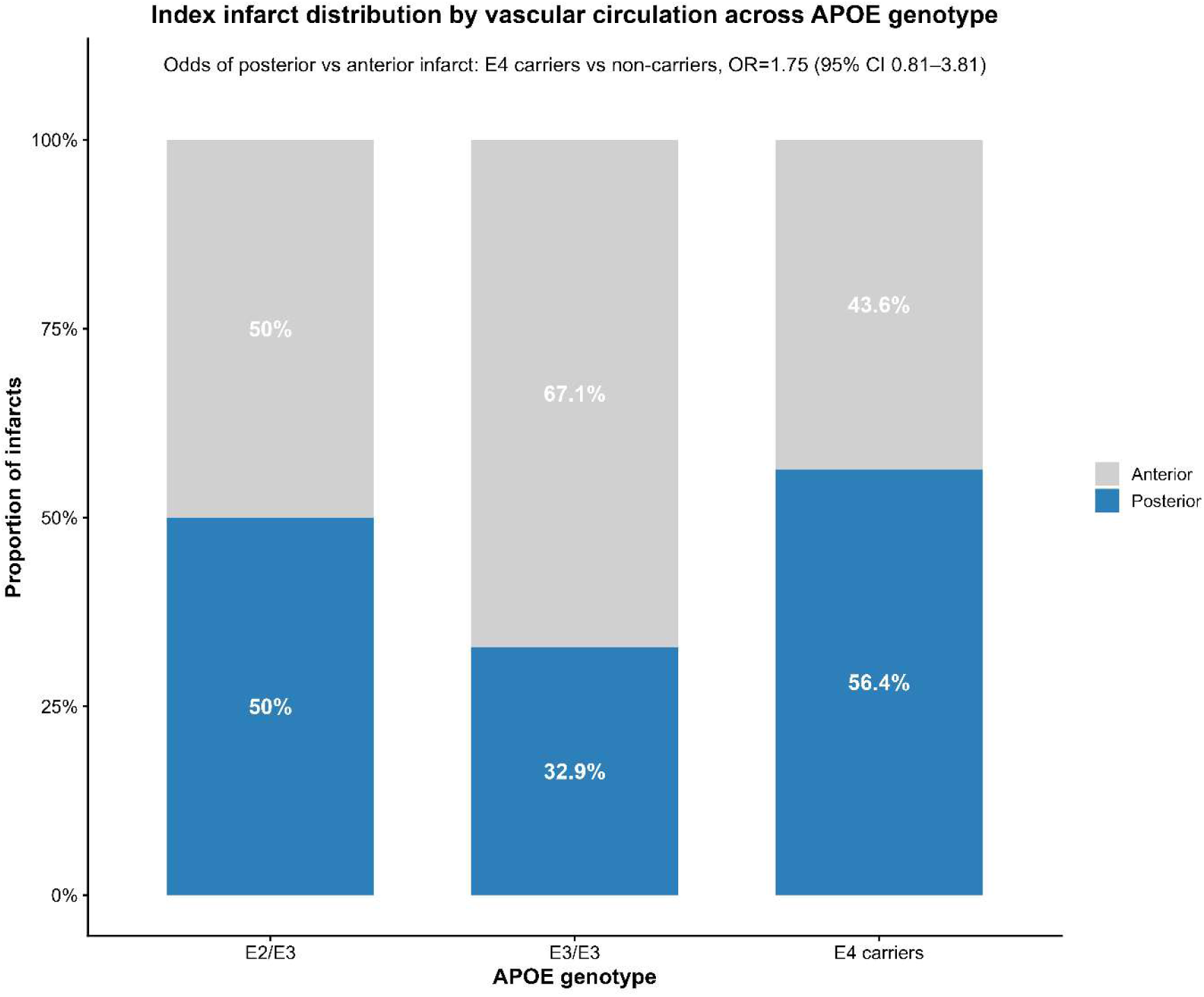
Distribution of index infarcts by vascular circulation and APOE genotype. Stacked bars show the proportion of infarcts in the anterior (grey) and posterior (blue) circulations within each genotype group. Percentages indicate the within-genotype distribution. The odds of a posterior versus anterior infarct were higher in APOE4 carriers compared with non-carriers (OR=1.75, 95% CI 0.81–3.81), adjusted for age and sex. Patients without visible infarcts were excluded from this analysis.

Binomial logistic regression, using a count-based model and adjusted for age and sex, indicated increased odds of posterior versus anterior circulation infarcts in E4 carriers compared with non-carriers (OR 1.75, 95% CI 0.81 to 3.81), although this finding did not reach statistical significance.

Anterior circulation infarcts (ACA cortex and anterior borderzone) were less frequent in E4 carriers (0% and 2.6%) than in other APOE groups (**Table 3**).

### 3.2 APOE genotype and BBB Metrics (PS, vP)

Overall, minimal variability was observed in PS and vP across genotype groups, and no statistically significant genotype-related differences were detected (**Figure 2; Supplementary Table 4**).

**Figure 2.**
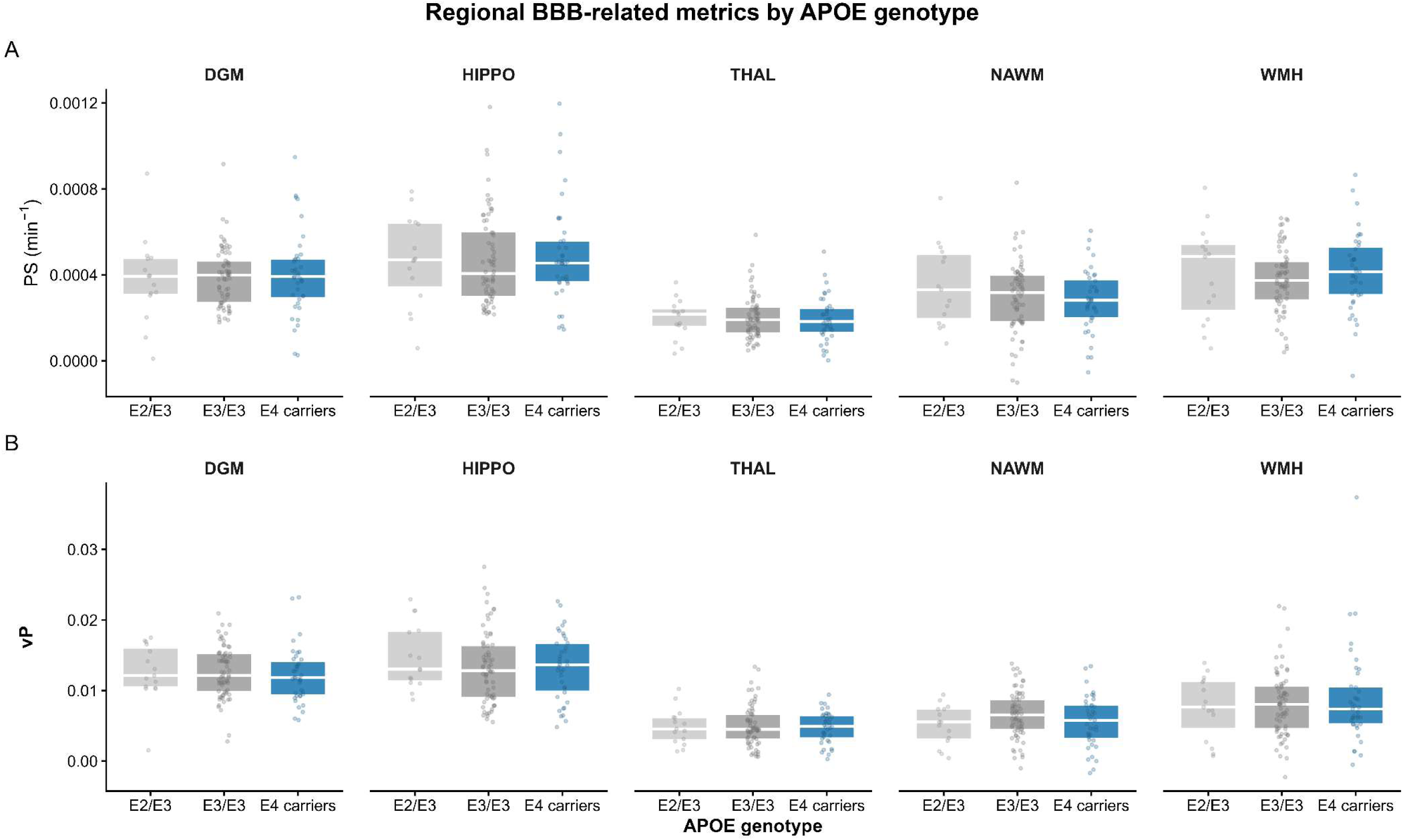
Regional BBB metrics (PS and vP) by APOE genotype. (A) Permeability–surface area product (PS, min⁻¹), reflecting BBB permeability. (B) Plasma volume fraction (vP), reflecting vascularity. Values are shown across deep grey matter (DGM), hippocampus (HIPPO), thalamus (THAL), normal-appearing white matter (NAWM), and white matter hyperintense regions (WMH), stratified by APOE genotype. Boxes indicate median and interquartile range; points represent individual patients. Plots show raw, unadjusted values. Covariate-adjusted associations accounting for age, sex, WMH volume, and vascular risk score are presented in Table 4.

**Table 4.**
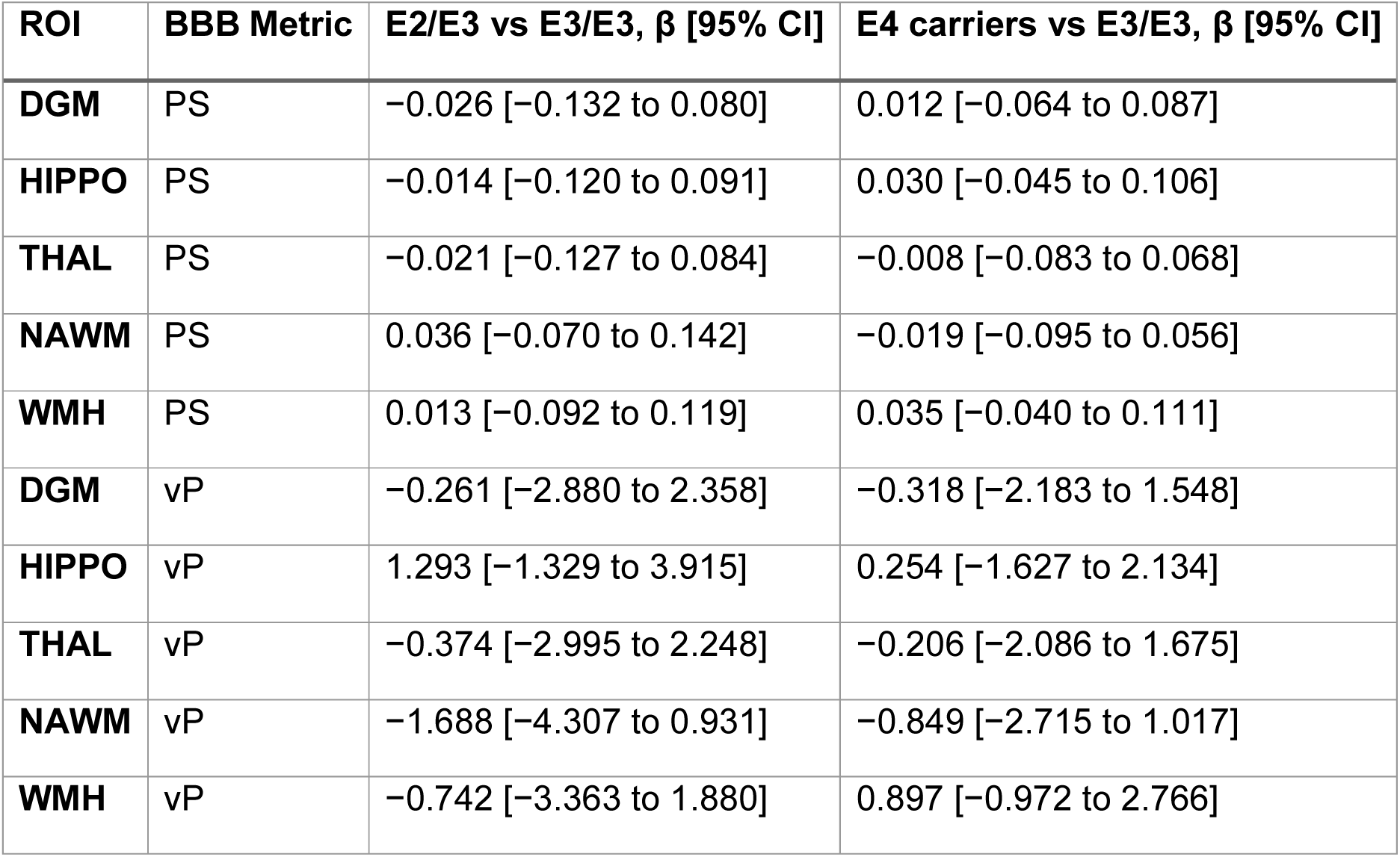
Associations between APOE genotype and blood–brain barrier metrics (PS and vP) across brain regions. Values are β estimates with 95% confidence intervals from linear mixed-effects models, including APOE genotype, ROI, age, sex, WMH volume, and vascular risk score (VRS) as fixed effects, with participant ID included as a random intercept. PS values (min⁻¹) are reported as ×10⁻⁴, and vP values (fractional plasma volume, unitless) as ×10⁻². DGM indicates deep grey matter; HIPPO, hippocampus; THAL, thalamus; NAWM, normal-appearing white matter; WMH, white matter hyperintensities.

| ROI | BBB Metric | E2/E3 vs E3/E3, $\beta$ [95% CI] | E4 carriers vs E3/E3, $\beta$ [95% CI] |
| --- | --- | --- | --- |
| DGM | PS | -0.026 [-0.132 to 0.080] | 0.012 [-0.064 to 0.087] |
| HIPPO | PS | -0.014 [-0.120 to 0.091] | 0.030 [-0.045 to 0.106] |
| THAL | PS | -0.021 [-0.127 to 0.084] | -0.008 [-0.083 to 0.068] |
| NAWM | PS | 0.036 [-0.070 to 0.142] | -0.019 [-0.095 to 0.056] |
| WMH | PS | 0.013 [-0.092 to 0.119] | 0.035 [-0.040 to 0.111] |
| DGM | vP | -0.261 [-2.880 to 2.358] | -0.318 [-2.183 to 1.548] |
| HIPPO | vP | 1.293 [-1.329 to 3.915] | 0.254 [-1.627 to 2.134] |
| THAL | vP | -0.374 [-2.995 to 2.248] | -0.206 [-2.086 to 1.675] |
| NAWM | vP | -1.688 [-4.307 to 0.931] | -0.849 [-2.715 to 1.017] |
| WMH | vP | -0.742 [-3.363 to 1.880] | 0.897 [-0.972 to 2.766] |

In linear mixed-effects models adjusting for age, sex, WMH volume, and VRS, APOE genotype was not associated with PS or vP across regions (**Table 4**).

Several covariate associations were observed (**Supplementary Table 5**). Increasing age was associated with lower PS in DGM, hippocampus, and thalamus, as well as lower PS and vP in WMH regions. Higher VRS was associated with lower vP in both NAWM and WMH regions.

In binomial logistic regression models adjusted for age, sex, and APOE4 carrier status, neither DGM plasma volume (vP; OR 1.28, 95% CI 0.43 to 3.99) nor DGM permeability (PS; OR 1.00, 95% CI 1.00 to 1.00) was significantly associated with posterior versus anterior circulation infarct presentation. Inclusion of DGM vascularity or permeability metrics did not substantively alter the association between APOE4 carrier status and infarct distribution. APOE4 effect estimates remained elevated across models. Increasing age was associated with reduced odds of posterior circulation infarcts (**Table 5**).

**Table 5.**
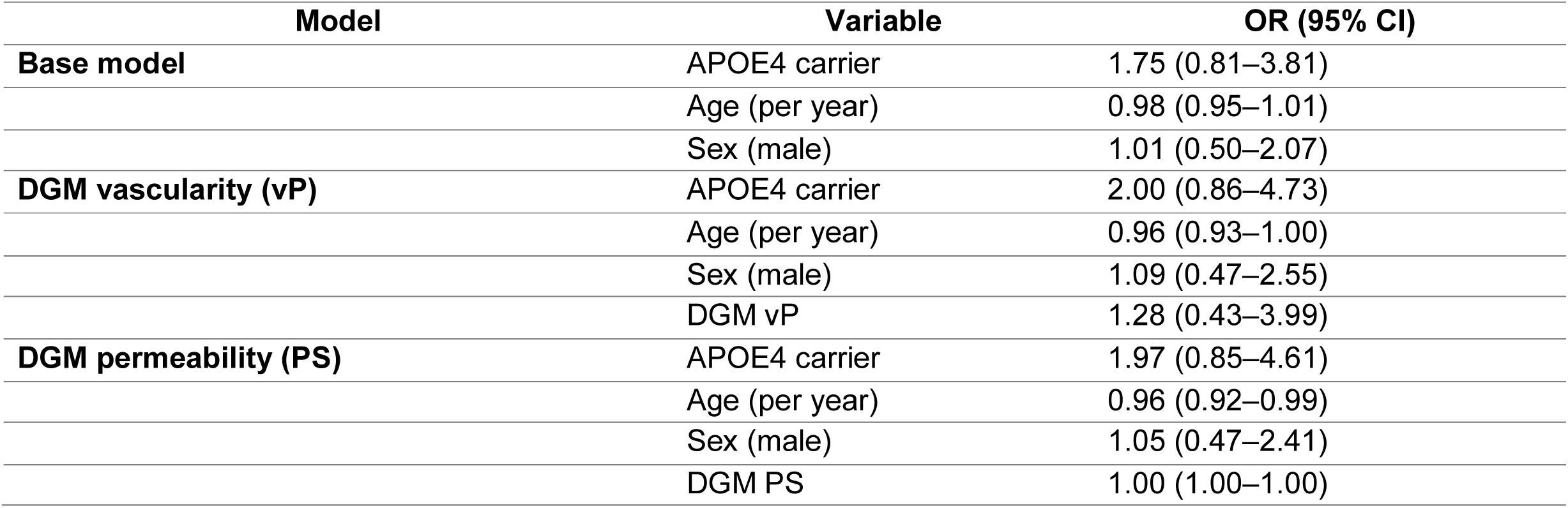
Logistic regression models for posterior versus anterior circulation infarcts. Values are odds ratios (ORs) with 95% confidence intervals from binomial logistic regression models assessing the odds of posterior versus anterior circulation infarcts. Models were adjusted for age and sex. DGM indicates deep grey matter; vP indicates vascularity (fractional plasma volume); PS, permeability–surface area product (BBB permeability).

## 4. Discussion

This study investigated APOE-related differences in infarct location, BBB permeability and plasma volume in patients with mild ischemic stroke. Quantitative DCE-MRI metrics and regional infarct mapping were used to assess whether APOE genotype influenced BBB permeability, regional vascularity, and infarct distribution. The principal findings were: (1) APOE4 carriers exhibited a trend toward a posterior-predominant infarct distribution albeit non-significant; (2) no genotype-related differences were observed in permeability–surface area product (PS) or plasma volume (vP), although age and vascular risk burden were associated with reduced PS and vP; and (3) there was a trend towards more BBB permeability and plasma volume in DGM in those with posterior rather than anterior location infarcts, again not significant. Collectively, these findings indicate that APOE4 genotype may influence infarct distribution, and associated microvascular dysfunction may manifest towards the posterior rather than anterior location.

Index infarct analysis revealed differences in infarct distribution across APOE genotypes, with E4 carriers demonstrating a non-significant trend toward a greater proportion of infarcts in posterior circulation territories. While E3/E3 individuals had a more diffuse distribution pattern across vascular territories, E4 carriers exhibited higher frequencies of PCA, posterior borderzone, and thalamic index lesions. Posterior circulation infarcts, supplied by the PCA, basilar artery, and intracranial and extracranial vertebral arteries, account for approximately 20% of all ischemic strokes and are commonly attributed to cSVD, LAA, or cardioembolism^55,56^. In this cohort, posteriorly located infarcts comprised 32.9% of infarcts in the E3/E3 group, 56.4% in E4 carriers, and 50% in the E2/E3 group.

This posteriorly weighted distribution is consistent with prior reports of WMH pathology in E4 carriers, which shows a predilection for parietal-posterior regions^13^. These regions are also commonly affected by cerebral amyloid angiopathy, an amyloid-driven disorder typically assessed by the presence of cerebral microbleeds on MRI. Lobar microbleeds frequently colocalise with WMH pathology, even in cognitively unimpaired individuals, suggesting shared regional vulnerability and potential overlap in underlying mechanisms^57^.

This pattern suggests a potential regional susceptibility associated with APOE4 carrier status to be confirmed in other studies; however, the underlying mechanism remains unclear. Further research is required to fully understand the underlying contributions of E4 carrier status to cerebrovascular health, particularly as it might pertain to a posteriorly weighted pattern of infarct involvement. Beyond supported research of APOE4’s role in atherosclerosis, exploratory research has begun to investigate the association between APOE4 and prothrombotic mechanisms at the cellular level and among cohort populations^58,59^. Considering APOE4’s association with mitochondrial energetics, endothelial function and prothrombotic factors, alterations in vascular stress or haemodynamic/platelet energetics, may represent potential mechanisms of interest.

Assessment of mean tracer kinetic parameters (vP and PS) showed a trend towards higher DGM PS and lower DGM vP among APOE4 carriers, but no statistically significant differences by APOE genotype, suggesting that DCE-MRI metrics may not be sufficient to differentiate genotype-related BBB pathophysiology at this cross-sectional time point (Table 4; Supplementary Table 4). When assessed in relation to infarct distribution, DGM permeability and vascularity were not significantly associated with posterior versus anterior circulation infarcts. However, inclusion of these metrics did not materially attenuate APOE4 effect estimates, suggesting that any relationship between APOE4 status and infarct topography is unlikely to be fully accounted for by regional DGM BBB characteristics measured in this study.

Interestingly, when vP and PS models were adjusted for WMH volume, VRS, and age, several non-genotype-related associations emerged in NAWM and WMH regions. In prior analysis of this cohort, PS decreased with age in DGM and WMH regions^21^, and in the present study, age-related reductions in PS were additionally observed in the hippocampus and thalamus. As discussed previously, the decline in PS is likely to reflect the decline in vascular surface area rather than a true decline in permeability but since the surface area is not known, this latter decline cannot be corrected for.

In WMH regions, reduced vP was also associated with elevated WMH volume and higher VRS (composite of hypertension, hyperlipidaemia, smoking history, and diabetes). Although no statistically significant genotype effects were observed, findings suggest a trend toward higher PS in DGM, hippocampal, and WMH regions among E4 carriers (Table 4).

Collectively, these findings are supportive of age-related influence on vascular pathophysiology in this stroke cohort. As PS reflects the product of vascular surface area and permeability, associations between tracer kinetic parameters and age in cSVD may be vulnerable to alterations in vessel density and size^21^. Such considerations may therefore affect interpretation, particularly in older patients with poorer vascular health, where reduced vascular surface area may be indicative of fewer vessels and, consequently, lower PS estimates.

Given the slightly younger age in the E4 carrier subgroup, and limited power in this study, these findings may warrant further investigation in a larger cohort. Despite the posterior trend observed in APOE4 carriers, no definite corresponding differences were observed in BBB permeability or vascularity, and regional BBB metrics were not associated with infarct localisation.

Associations of APOE4 with ischemic stroke remain controversial, largely due to inconsistent evidence regarding APOE4’s role as a genetic risk factor, and whether it acts as a contributing, compounding, or conflicting factor in IS pathophysiology^2,6–9,60^. Meta-analytic evidence^10^ suggests that E4 increases cSVD risk but has no significant effect on LAA, underscoring the complexity of APOE’s relationship with IS^61^.

## Limitations

Several limitations should be considered when interpreting these findings. First, relatively modest sample size limited statistical power to detect small genotype-related effects, particularly in subgroup analyses involving APOE4 carriers and infarct location. Although APOE genotype frequencies in our cohort were comparable to other predominantly European-ancestry datasets (UK Biobank^51^, Lothian Birth Cohort^52^, and CHARGE^53^), the overall number of participants with both APOE and DCE-MRI data was limited.

Second, methodological considerations related to BBB permeability measurement using DCE-MRI should be acknowledged. Interpretation of PS is dependent on both vascular permeability and vascular surface area, which cannot be independently quantified using the present approach. Consequently, age- and cSVD-related reductions in vascular density may influence PS estimates and their interpretation.

Finally, the limited ethnic diversity of the cohort restricts the generalisability of these findings. Given that APOE4-associated risk affects carriers across populations, broader representation should be prioritised in future studies.

## Disclosures

### Data Availability

data are available from the senior corresponding author upon reasonable request.

## Non-standard Abbreviations and Acronyms

AD: Alzheimer’s disease
APOE: apolipoprotein
E BBB: blood–brain barrier
CE: cardioembolic
CI: confidence interval
cSVD: cerebral small vessel disease
DCE-MRI: dynamic contrast-enhanced magnetic resonance imaging
DTI: diffusion tensor imaging
FLAIR: fluid-attenuated inversion recovery
FSPGR: fast-spoiled gradient-echo
DGM: deep grey matter
ICV: intracranial volume IS ischemic stroke
LAA: large artery atherosclerotic
mRS: modified Rankin Scale
NIHSS: NIH Stroke Scale
OR: odds ratio
PS: permeability–surface area product
PVS: perivascular spaces
ROI: region of interest
VaD: vascular dementia
VIF: vascular input function
vP: fractional plasma volume
VRS: vascular risk score
WMH: white matter hyperintensity

## 5. Acknowledgements

The authors thank the imaging technologists, data managers, and clinical staff involved in the Mild Stroke Study 2 (MSS2) for their assistance with patient recruitment and MRI acquisition. The authors are also grateful to all participants of the MSS2 for their time and commitment to the study.

## 7. Disclosures

### 8. Author Contributions

Krystal Laing conceived and designed the study, performed the analyses, interpreted the data, and drafted the manuscript. Francesca M. Chappell advised on statistical modelling and provided Mild Stroke Study 2 (MSS2) data. Michael J. Thrippleton provided expertise on Patlak modelling and the derivation of DCE-MRI parameters. Maria del C. Valdés-Hernández conducted the image processing in the primary study, oversaw MRI segmentation for hippocampal and thalamic signal enhancement and contributed to generation of signal enhancement curves. Owen Dando and Deepali Vasoya performed APOE genotype imputation. Stephen Makin recruited the MSS2 patients and collected the clinical data. Paul Armitage designed and implemented the MRI acquisition protocol. Joanna M. Wardlaw designed the study, obtained funding, oversaw the study conduct and analysis, interpreted the data, drafted and edited the paper, and is the study guarantor and principal investigator. All authors critically revised the manuscript and approved the final version.

